# Development of a flexible split prime editor using truncated reverse transcriptase

**DOI:** 10.1101/2021.08.26.457801

**Authors:** Chunwei Zheng, Shun-Qing Liang, Bin Liu, Pengpeng Liu, Suet-Yan Kwan, Scot A. Wolfe, Wen Xue

## Abstract

Prime Editor (PE) has tremendous promise for gene therapy. However, it remains a challenge to deliver PE (>6.3 kb) *in vivo*. Although PE can be split into two fragments and delivered using dual adeno-associated viruses (AAVs), choice of split sites within Cas9 – which affects editing efficiency – is limited due to the large size of PE. Furthermore, the potential effect of overexpressing RT in mammalian cells is largely unknown. Here, we developed a compact PE with complete deletion of the RNase H domain of reverse transcriptase (RT), which showed comparable editing to full-length PE. Using compact PE, we tested the effect of 4 different Cas9 split sites and found that the Glu 573 split site supports robust editing (up to 93% of full-length PE). The compact PE, but not PE2, abolished its binding to eRF1 and showed minimal effect on stop codon readthrough, which therefore might reduce the effects on protein biosynthesis. This study identifies a safe and efficient compact PE2 that enables flexible split-PE design to facilitate efficient delivery *in vivo* and advance the utility of prime editing.

## Introduction

Prime editor, a new gene editing tool, consists of a Cas9 nickase fused to an engineered Moloney murine leukemia virus (MoMLV) RT. In the PE construct, Cas9 nickase is guided by RNA (sgRNA) to a target site, where the nickase cleaves only one DNA strand. MoMLV RT then uses an extension of the sgRNA, called prime editing guide RNA (pegRNA), as an RT template to generate complementary DNA for precise repair of the nicked site. Using PE, all possible base substitutions, small insertions (up to 44 bp), and small deletions (up to 80 bp) can be flexibly generated^1,2^, which enhances the versatility and precision of CRISPR gene editing.

For PE to succeed as a tool for gene therapy, it is also important to deliver PE efficiently *in vivo*. Adeno-associated virus (AAV) has emerged as the most widely used vector for *in vivo* delivery of gene therapies. However, the packaging capacity of AAV vectors is limited to ~5 kb^3,4^, which is insufficient for PE (>6.3 kb). To address this issue, Cas9 is split into two vectors and delivered by dual AAVs. Among strategies to split Cas9^5–8^, split-intein systems are the most extensively studied. Different split sites in Cas9 can affect its reconstituting activity and structural stability, which results in different gene editing efficiency. However, due to the large size of PE, split sites in Cas9 of PE are limited to a small region to fit within AAV vectors, limiting prime editing *in vivo*.

Furthermore, the effect of overexpressing RT required comprehensively analysis for their utility. Previous studies showed MoMLV RT interacts with eRF1 through RNase H domain^9^. eRF1 plays an essential role in termination of protein translation and nonsense-mediated decay (NMD) of mRNA molecules^10^. During regular translational termination, eRF1 recognizes all 3 stop codons and recruits other termination factors such as eRF3 to form the pre-termination complex. However, when eRF1 detects a premature stop codon or an irregularly spliced mRNA, it will recruit upframeshift protein 1 (UPF1) to trigger NMD^10^. eRF1 interacts with RT and other regulatory factors through its C-terminal domain. RT may compete with eRF3, UPF1 and other regulatory factors’ binding to eRF1, thereby promoting stop codon readthrough and bypassing mRNA degradation by NMD. Although overexpressing PE2 showed minimal perturbation of the transcriptome relative to Cas9 nickase or inactivated RT, the effect of PE2 overexpression on protein translation has not been investigated.

In this study, we developed a more compact PE (cPE2) with flexible split sites in Cas9. The cPE showed comparable editing efficiency in HEK293 cells. Using the cPE2, we developed an optimized split-PE to support robust prime editing in HEK293T cells and mouse liver. We also showed overexpressed PE2 interacted with eRF1 through RNase H domain to promote readthrough in HEK293T cells, while cPE2 without RNase H domain abolished the interaction and the readthrough effect. This cPE has the potential to improve the safety and delivery efficiency of prime editing for *in vivo* applications, including gene therapies.

## Results

### Development of a compact PE lacking the RNase H domain

MoMLV RT is a monomeric polypeptide containing fingers/palm, thumb, connection, and RNase H domains (residues 1-275, 276-361, 362-496, 497-671, respectively)^11–13^. It has two active sites (DNA polymerase and RNase H) that reside in separate, distinct structural domains^14^ (**Fig. 1a**). Mutating the RNase H domain, which is responsible for degrading RNA in the RNA/DNA hybrid during DNA synthesis, can improve the efficiency of cDNA synthesis from the protected RNA^15^. We hypothesized, therefore, that the RNase H domain is not essential for prime editing. To this end, we constructed four compact PE2 variants – RT497, RT474, RT418, and RT362 – with progressive truncation of the RNase H and connection domains **(Fig. 1a)**. After transfection into HEK293T cells, we observed the successful expression of each PE2 variant by western blot (**Supplementary Fig. 1a**).

**Fig. 1:**
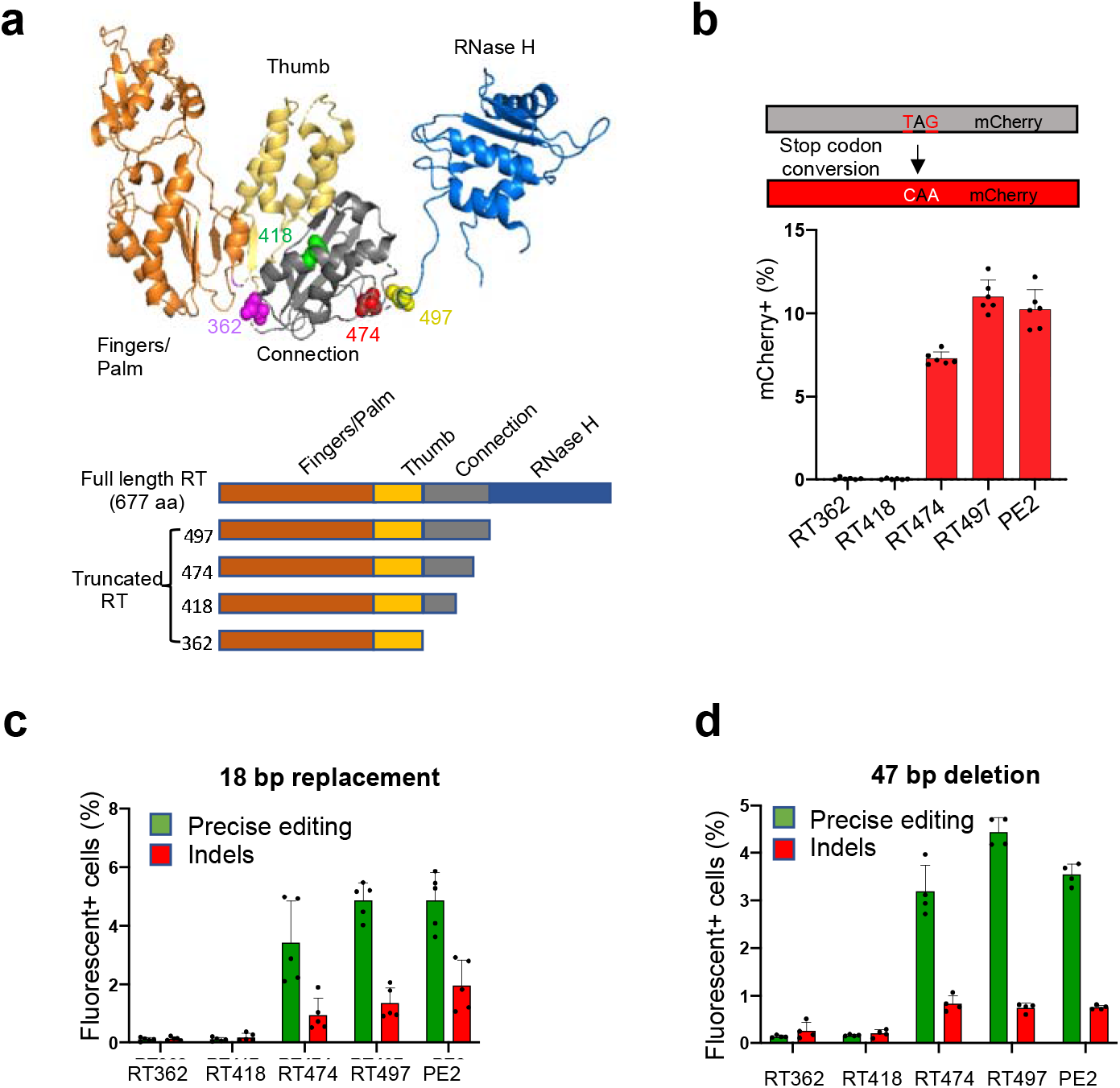
The RT RNase H domain in PE2 is dispensable for prime editing. **a** The crystal structure of MoMLV RT (top, PDB: 5dmq.1)^9^. Schematic representations of RT variants with C-terminal truncations (bottom). **b** Schematic of an mCherry reporter containing a premature stop codon in HEK293T cells. Prime editors can generate a TAG-to-CAA transition to correct the premature stop codon (top). Editing efficiencies of PE2 variants (RT362, RT418, RT474, and RT497) and full-length PE2 were measured by flow cytometry (bottom). Dashed line denotes PE2 levels. Results were obtained from six independent experiments, shown as mean ± s.d. **c** Traffic light reporter Multi-Cas Variant 1 (TLR-MCV1) HEK293T cells containing a GFP with a 39-bp insertion, P2A, and out-of-frame mCherry. A precise insertion of 18-bp to replace the 39-bp insertion leads to GFP expression, while indels with a frameshift lead to mCherry expression. The frequencies of precise insertion (GFP+) or indel (mCherry+) were measured by flow cytometry. Results were obtained from five independent experiments, shown as mean ± s.d. **d** TLR HEK293T cells containing a GFP with 47-bp insertion. The frequencies of precise deletion of 47-bp (GFP+) or indel (mCherry+) were measured by flow cytometry. Results were obtained from four independent experiments, shown as mean ± s.d.

To evaluate the editing efficiency of the four PE2 variants in HEK293T cells, we used an mCherry reporter cell line with a premature TAG stop codon in the mCherry coding sequence (**Fig. 1b**). We found that RT497, in which the RNase H domain is completely removed, exhibited prime editing efficiency that was similar to PE2 (11.0% vs 10.2%) (**Fig. 1b, Supplementary Fig. 1b**). By contrast, RT474, RT418, and RT362, which lack not only the RNase H domain but also the connection domain, showed decreased editing efficiencies (7.3%, <0.1%, and <0.1%, respectively).

Next, we tested the efficiency of precise insertion of PE2 variants at a target site using the Traffic Light Reporter Multi-Cas Variant 1 (TLR-MCV1) HEK293T reporter line for testing an 18-bp insertion (**Supplementary Fig. 2a)**^16,17^. Consistent with our results in mCherry reporter cells, RT362, RT418 and RT474 showed decreased on-target editing efficiencies (0.1%, 0.1%, and 3.4%, respectively). By contrast, RT497 exhibited an editing efficiency that was similar to that of PE2 editing efficiency (4.9% for both). The indel rate of RT497 was also similar to PE2 (1.3% vs 2.0%, **Fig. 1c**).

Finally, we tested the efficiency of precise deletion of PE2 variants using a second TLR reporter for testing a 47-bp insertion (**Supplementary Fig. 2b)**^16^. We found that RT474 and RT497 exhibited editing efficiencies that were similar to PE2 (3.2%, 4.4%, and 3.5%, respectively), while the editing efficiencies of RT362 and RT418 were decreased (0.1% and 0.2%) (**Fig. 1d**).

Collectively, these results demonstrate that the RT RNase H domain is dispensable for prime editing, and that RT497 exhibits nucleotide substitutions, precise small insertions and deletions, and indel rates at comparable efficiencies to that of full-length PE2.

### Compact PE efficiently introduces mutations at endogenous loci

To show that the observed editing efficiencies of PE2 variants are not specific to fluorescent reporters, we further tested PE2 variant editing at the +1 position (1-bp after the Cas9 nicking site) in four previously-validated endogenous loci (*FANCF*, *VEGFA*, *RNF2*, *HEK3*) by amplicon sequencing^1^. Consistent with our results using fluorescence reporter systems, RT497 induced comparable on-target editing efficiencies as PE2 at all four loci. Indeed, RT497 induced a 1-bp substitution at a frequency of 51.1%, 50.7%, and 20.7% at *FANCF*, *VEGFA*, and *RNF2*, respectively. PE2 resulted in 1-bp substitutions at a frequency of 46.6%, 46.9%, and 20.3% at *FANCF*, *VEGFA*, and *RNF2*, respectively. RT497 and PE2 induced 3-bp insertions at a frequency of 40.9% and 37.7%, respectively, in the *HEK3* locus (**Fig. 2a, Supplementary Fig. 3a-d**). RT474 exhibited lower editing efficiency than PE2 for 1-bp substitutions at *RNF2* (12.6% vs 20.3%), but comparable editing efficiencies for 1-bp substitutions at *FANCF* (46.7% vs 46.6%) and *VEGFA* (48.9% vs 46.9%), and 3-bp insertion at *HEK3* (37.8% vs 37.7%).

**Fig. 2:**
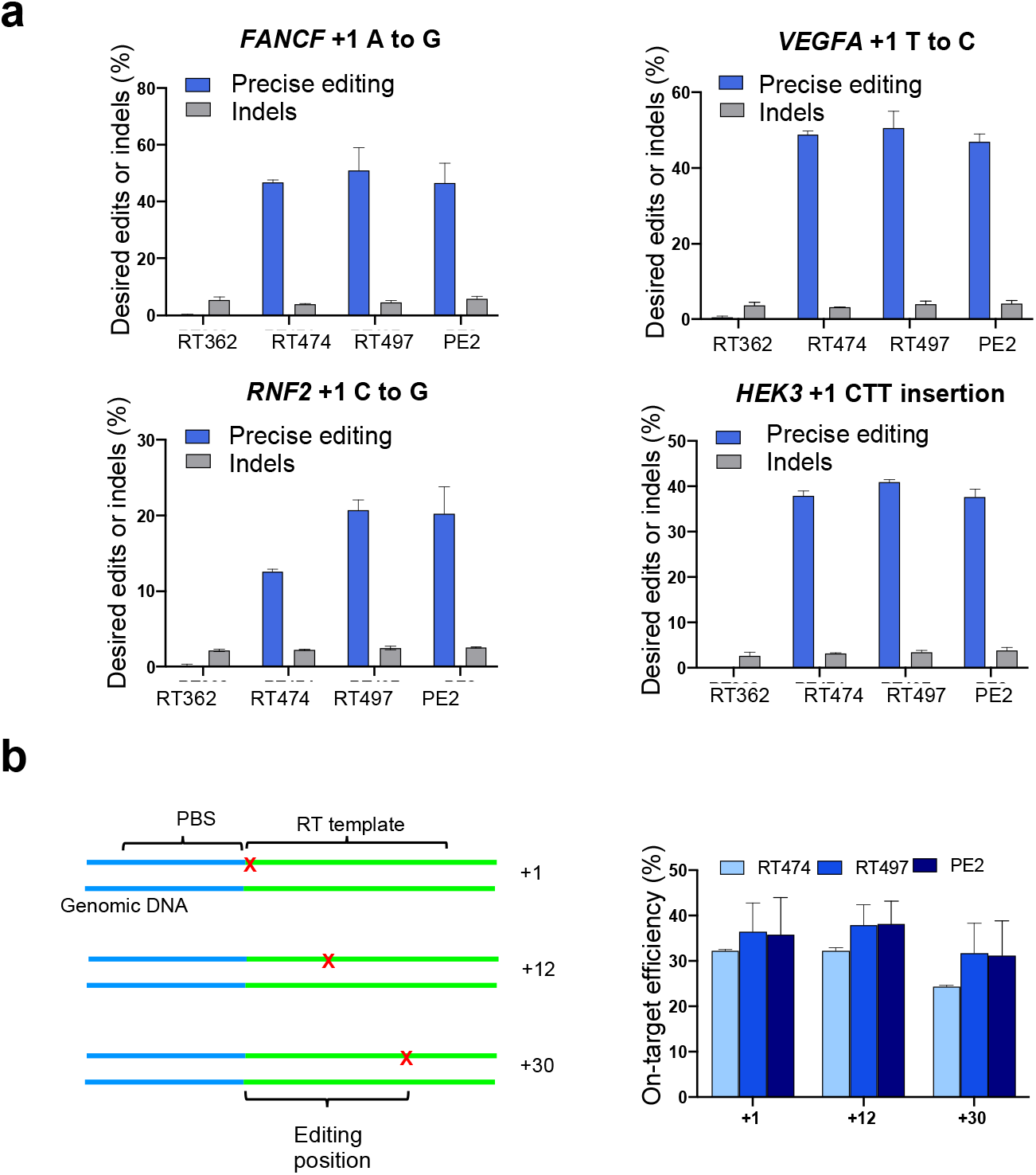
Editing frequency of truncated prime editors at endogenous loci. **a** Comparison of precise editing and indel efficiency for substitutions and insertions induced by RT362, RT474, RT497, and PE2 at *FANCF*, *VEGFA*, *RNF2*, *HEK3* loci in HEK293T cells. **b** Schematic representation of single nucleotide substitution at positions +1, +12, +30 of *HEK3* (left). Comparison of precise editing efficiency for single nucleotide substitution at positions +1, +12, +30 of *HEK3* by RT474, RT497 and PE2 (right). Results were obtained from three independent experiments (RT474) or four independent experiments (RT362, RT497, PE2), shown as mean ± s.d.

Although the RNase H domain of MoMLV RT is not required for DNA polymerase activity, the residues in the C helix may stabilize enzyme/primer-template interactions. Thus, loss of the RNase H domain could affect the processivity of DNA synthesis, leading to a shorter RT product^18^. To determine whether RT497 have a narrower editing window compared to PE2, we compared the editing efficiencies of each (1-bp substitutions) at positions +1, +12 and +30 in the *HEK3* locus (**Fig. 2b**)^1^. RT497 and PE2 had similar editing efficiencies at +1 position (36.5% vs 35.8%), +12 position (38.0% % vs 38.2%), and +30 position (31.7% vs 31.2%). However, RT474 showed a slightly lower editing efficiency at the +1 position (32.3% vs 35.8%), +12 position (32.3% to 38.2%), and +30 position (24.4% to 31.2%) (**Fig. 2b**). These data suggest that truncating the RNase H domain in MoMLV RT does not affect the processivity of RT for short products (~30 bp). Based on these results, we elected to move forward with RT497 (cPE2) to develop an efficient split-PE.

### Development of efficient split-PE

Due to the packaging limit of AAV vector (<5 kb) and the large size of SpyCas9 (4.1 kb) and MoMLV RT (2 kb), Cas9 could only be split after residue R691 to package the C-terminal fragment of PE2 with U1a promote, RBG poly A, and *Nostoc punctiforme* (Npu) intein into one AAV vector. Here, our newly engineered cPE2 confers flexibility in the choice of split site. The smaller size of cPE2 allows Cas9 to be split up to residue L551 from the C-terminal (**Supplementary Fig. 4a**), enabling the use of the potentially more efficient E573 Cas9 split site^3,19^. To identify an optimal split cPE2, we generated 4 different split sites that have been previously tested for split-Cas9 or split-base editors: E573 (split-cPE2-573), K637 (split-cPE2-637), Q674 (split-cPE2-674), or Q713 (split-cPE2-713)^3,8,19,20^. To determine the effect of each split site on cPE2 fusion, we co-transfected the N-terminal and C-terminal fragments of split-PE2 variants into HEK293T cells and measured protein expression using western blot. We observed spliced full-length cPE2 expression after transfection of split-cPE2-573, split-cPE2-674, and split-cPE2-713, whereas only unspliced cPE2 was detected after transfection of split-cPE2-637. Among them, split-cPE2-573 showed the highest full-length expression level (**Fig. 3a**).

**Fig. 3:**
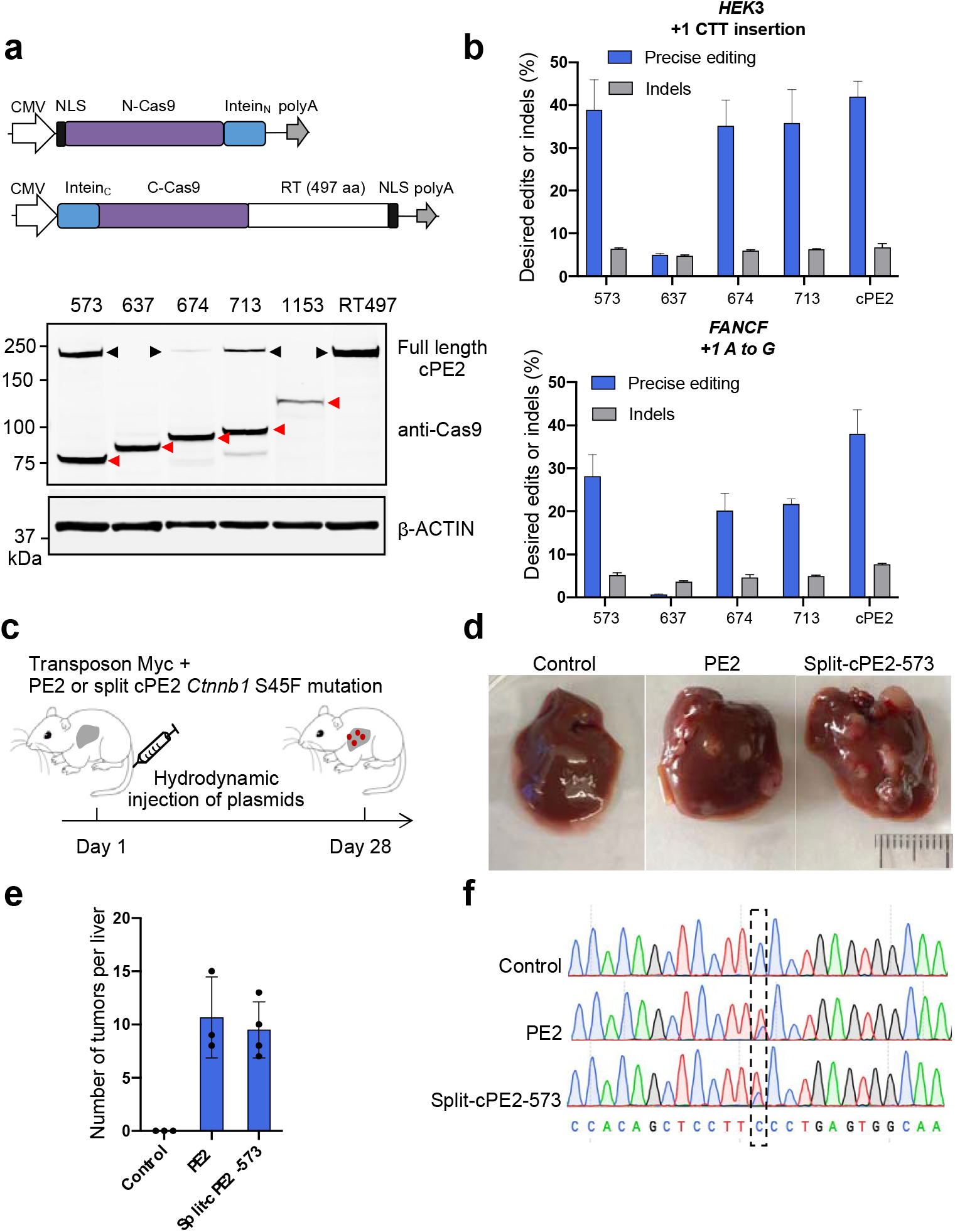
Development of a split-intein cPE2 for *in vivo* delivery. **a** Map of split-cPE2 plasmids (top). CMV, cytomegalovirus promoter; NLS, SV40 nuclear localization sequence; N-Cas9, N-terminal Cas9 nickase; C-Cas9, C-terminal Cas9 nickase; poly A, bGH poly A (top). Western blot showing split-cPE2 and unsplit cPE2 expression (bottom). N-terminal cPE2 and C-terminal cPE2 of each variant were co-transfected into HEK293T cells. As a postive control, unsplit cPE2 plasmid was transfected. Cell lysates were probed with anti-GAPDH and anti-Cas9. Unspliced (red arrows) and reconstituted cPE2 (black arrows) were detected using anti-Cas9 antibody. β-actin was used as a loading control. Unspliced and reconstituted cPE2 were observed after transfection of split-cPE2-573, split-cPE2-674, and split-cPE2-713 (lanes 1,3,4). Only unspliced cPE2 (red arrows) was detected after transfection of split-cPE2-637 and split-cPE2-1153 (lanes 2,5). **b** Editing frequencies of split-cPE2 at *FANCF* and *HEK3* in HEK293T cells. Results were obtained from three independent experiments, shown as mean ± s.d. **c** PE2 or split-cPE2-573, pegRNA, nicking sgRNA targeting Ctnnb1, MYC transposon, and transposase plasmids were hydrodynamically injected into wild-type FVB mice to generate liver tumor. PE2, non-target pegRNA, MYC transposon and transposase plasmids were injected in the control group. **d** Representative images of liver tumor burden after plasmid injection. **e** Mean number of visible tumor nodules were counted in mouse livers 28 days after plasmids injection. Results were obtained from three (control and PE2) or four (split-cPE2-573) independent experiments, shown as mean ± s.d. **f** Sanger sequencing of control liver and representative tumors.

Next, we tested the prime editing efficiency of each split-cPE2 at two endogenous loci (*FANCF* and *HEK3*). Using deep sequencing, we found that split-cPE2-573 showed the highest precise editing efficiency among split-cPE2 variants. Split-cPE2-573 editing efficiency was 92.6% of full-length cPE2 at *HEK3* (38.9% vs 42.0%), and 74.2% of cPE2 at *FANCF* (28.2% vs 38.0%) (**Fig. 3b, Supplementary Fig. 4b**). In a previous study, PE2 had been split at 1005 and 1024 with *Rhodothermus marinus* (Rma) intein^21^, and these two split-PE2 showed ~72.0% and ~75.8% editing efficiency compare with full-length PE2 at *HEK3* loci, ~65.6% and 68.4% editing efficiency compare with full-length PE2 at *VEGFA* loci^21^, while the Split-cPE2-573 showed higher editing efficiency, ~92.6% and ~78.5% of full-length cPE2 at *HEK3* and *VEGFA* loci, respectively (**Fig. 3b, Supplementary Fig. 5**).

Finally, we sought to determine whether split-cPE2-573 supports *in vivo* delivery of functional prime editor. We designed a pegRNA to generate a C:G to T:A transversion in *Ctnnb1* (β-Catenin) that produces the oncogenic S45F mutation frequently observed in liver cancer^22,23^. PE2 or split-cPE2-573, and pegRNA were delivered to the liver of FVB mice via hydrodynamic tail vein injection. MYC transposon and transposase were co-injected to provide a second oncogenic driver necessary for liver cancer formation with the *Ctnnb1* mutation (**Fig. 3c**)^24^. Mice injected with PE2 (n=3) and split-cPE2-573 (n=4) developed a similar number of liver tumor nodules after 28 days, with an average of 10.7 tumors and 9.5 tumors per mouse respectively (**Fig. 3d, e**). Sanger sequencing of tumor nodules showed precise substitution of S45F in *Ctnnb1* in both groups (**Fig. 3f**). As a control, a non-target pegRNA was injected and no tumor was formed. Together, these data demonstrate that split-cPE2-573 can be used for gene editing *in vivo*. Using split-cPE2-573, the coding sequence of PE2 could be distributed on a dual-vector system and packaged into AAV.

### PE2 but not cPE2 promotes stop codon readthrough

Previous crystal structure studies revealed that MoMLV RT interacts with the C-terminal domain of eukaryotic translation termination factor 1 (eRF1) via the RNase H domain (**Fig. 4a**)^9^. As PE2 fused with MoMLV RT with D200N, L603W, T330P, T306K and W313F mutation, we tested whether PE2 or cPE2 interacts with eRF1. Co-immunoprecipitation assays showed that eRF1 interacted with full-length PE2, but not cPE2 (**Fig. 4b**), probably because cPE2 fused with truncated MoMLV RT without RNase H domain. Structure of eRF1/eRF3 and MoMLV RT/eRF1 complexes showed that eRF1 interacted with both MoMLV RT and eRF3 through overlapping surface regions^9^, therefore, RT likely outcompetes eRF3 binding to eRF1. To test whether PE2 or cPE2 could promote readthrough, we used an established stop codon readthrough reporter containing *Renilla* luciferase, a recoding window that includes a stop codon, and firefly luciferase (*Renilla*:TAG:firefly)^25^ **(Fig. 4c)**. Stop codon readthrough is measured by firefly relative to *Renilla* luciferase activities. As a control, we used a reporter containing a recoding window lacking a stop codon (*Renilla*:CAG:firefly). PE2 overexpression led to a ~1.6 fold (4.6% vs 7.6%) increases in readthrough, while cPE2 overexpression showed comparable readthrough rates as the control construct **(Fig. 4d).** Taken together, these data suggest PE2 interacted with eRF1 via the RT RNase H domain, thereby promoting stop codon readthrough **(Fig. 4e)**.

**Fig. 4:**
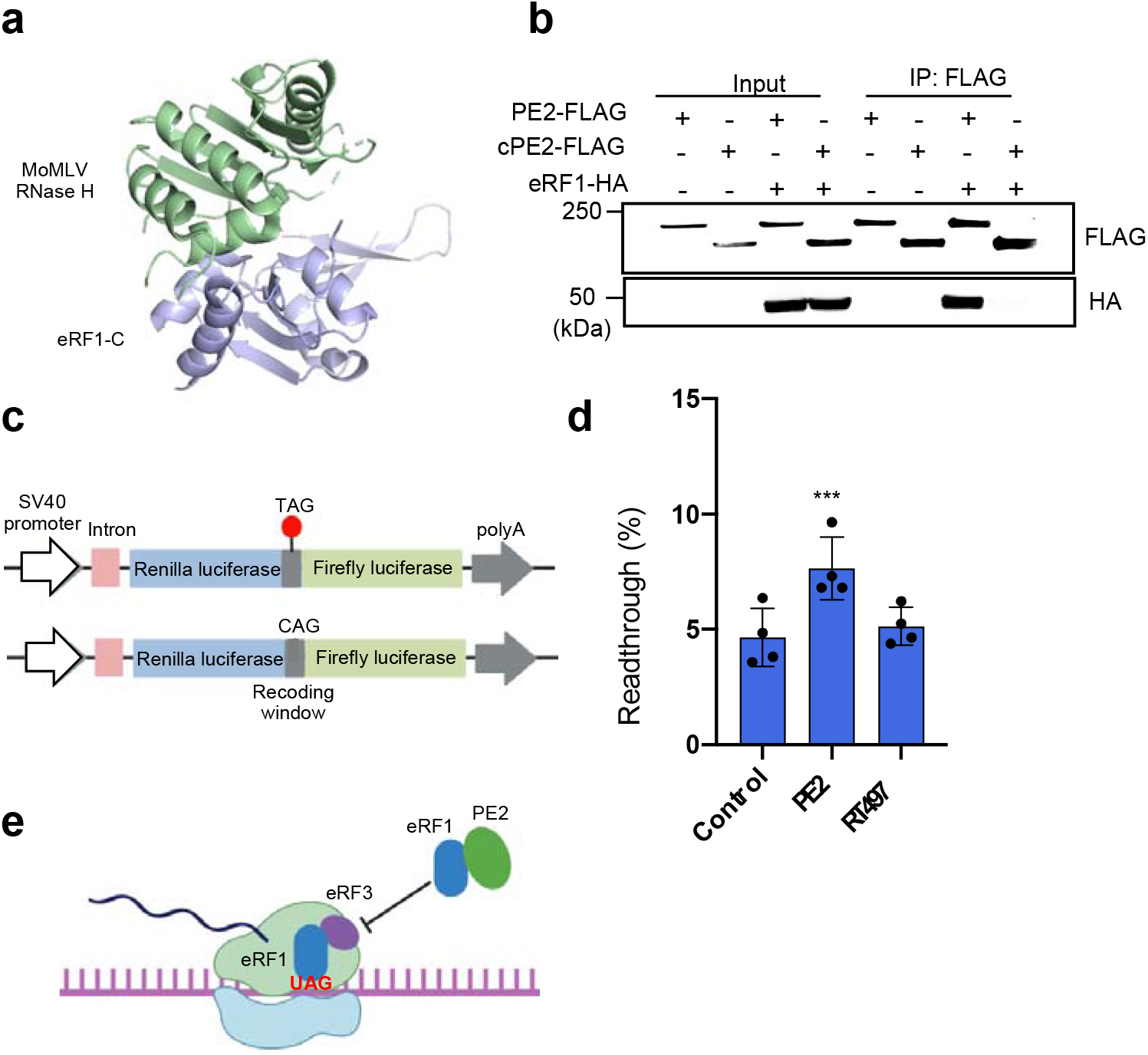
PE2 increases stop codon readthrough through the RNase H domain. **a** The structure of MoMLV RNase H/C-eRF1 complex (PDB: 5DMQ)^9^. **b** Co-immunoprecipitation assays for interactions of PE2 with eRF1. Plasmid construct of PE2-FLAG or cPE2-FLAG were co-transfected with eRF1-HA into HEK293T cells. Empty vectors expressing each tag (FLAG, HA) were used as negative controls. **c** Schematic of the dual luciferase bicistronic vector used in translational readthrough assays. The red dot represents the stop codon. **d** PE2 but not cPE2 overexpression promotes stop codon readthrough. PE2, cPE2 or control vector were co-transfected with dual luciferase reporter in HEK293T cells. The ratio of Firefly:Renilla from each experimental was normalized to a control group reporter lacking a stop codon between Firefly and Renilla. **e** Model of PE2 overexpressed promoting stop codon readthrough. PE2 interacts with eRF1 via the RNase H domain, which inhibits the binding of eRF1 with eRF3, promoting stop codon readthrough.

## Discussion

Prime editors have the potential to correct human pathogenic mutations by generating precise transitions, transversions, small insertions, or small deletions. However, the large size of prime editors makes them more difficult to deliver *in vivo* than Cas9 or base editors - a limitation impeding the developing of PE-based gene therapies. Recently, *in vivo* delivery of prime editors using dual-AAVs has been reported^16,21^. However, these studies used 713 split site in Cas9 with strict limitations in promoter and poly A length^16^, or used suboptimal split Cas9 sites 1005 and 1024^21^. In these cases, prime editing efficiency was suboptimal *in vivo*.

Expression of PEs perturbs the transcriptome minimally compared to expression of Cas9 and RT^1^. However, previous studies reported that overexpressing MoMLV RT could interact with eRF1 via RNase H domain to occlude the binding of eRF3 to eRF1, thereby promoting stop codon readthrough^9,26^. Hence, overexpressed PE2 may affect protein biosynthesis through binding to eRF1. Given that PEs maybe used for gene therapy in the future, investigations into their potential effects of protein translation are warranted.

In this work, we report that the MoMLV RT RNase H domain is dispensable for prime editing. By removing the RNase H domain, we developed a compact PE2 (cPE2) with comparable editing and indel efficiencies to full length PE2. we developed a compact PE with comparable editing efficiency to full length PE. Using the compact PE, we screened 4 split sites combined with Npu intein. Consistent with previous studies using base editors, we found split-cPE2-573 showed the highest on-target editing efficiency among split sites at endogenous loci^3,8,19^. Furthermore, we also found the compact PE without RNase H domain has no interaction with eRF1 which maintains the basal level of stop codon readthrough.

In summary, this study suggests that the RNase H domain of RT is dispensable for prime editing. PE2 could interact with eRF1 via RNase H domain thereby promote stop codon readthrough. The new version of compact PE2 enables flexible split site design to facilitate *in vivo* delivery.

## Material and methods

### Plasmid construction

Plasmids expressing sgRNA were generated by ligation of annealed oligos into a BfuA I-digested vector ^16^. For plasmids expressing plasmids, gBlocks gene fragments were inserted into a BfuA I/EcoR I-digested vector by Gibson assembly ^16^. Compact PE2 variants were generated by PCR using Phusion master mix (ThermoFisher Scientifc). Sequences of sgRNA and pegRNA are listed in Supplementary Table 1. All primers used for cloning are listed in Supplementary Table 2. All plasmids used for mammalian cell experiments were purified using plasmid Miniprep kits (Qiagen).

### Cell culture

Human embryonic kidney (HEK293T) cells were purchased from ATCC and cultured in Dulbecco’s Modified Eagle’s Medium (DMEM) supplemented with 10% (v/v) fetal bovine serum (Gibco) and 1% (v/v) Penicillin/Streptomycin (Gibco). Cells were cultured at 37°C with 5% CO_2_ and tested negative for mycoplasma.

### HEK293T transfection and genomic DNA extraction

HEK293T and HEK293T reporter cells were seeded on 12-well plates at 100,000 cells per well. 24 hours after seeding, cells were transfected using Lipofectamine 3000 reagent (Invitrogen) following the manufacturer’s protocol. Briefly, 1 μg PE2, 330 ng pegRNA, and 110 ng nicking sgRNA were transfected using 3 μL Lipofectamine 3000 and 3 μL P3000 (2 μl/μg DNA). For split-cPE2, HEK293T cells were co-transfected with 1 μg N-terminal fragment, 1 μg C-terminal fragment, 330 ng pegRNA, and 110 ng nicking sgRNA using 3 μL Lipofectamine 3000 and 5 μL P3000. For genomic DNA extraction from cells, cells were cultured for 3 days after transfection, washed with PBS, pelleted, lysed with 100 μL Quick extraction buffer (Epicenter), and incubated at 65°C for 15 min and 98°C for 5 min. To extract genomic DNA from mouse liver tissue, PureLink Genomic DNA Mini Kit (Thermo Fisher) was used following the manufacturer’s protocol.

### Co-immunoprecipitation assay

Plasmid construct of PE2-FLAG or RT497-FLAG were co-transfected with eRF1-HA into HEK293T cells. Empty vectors were used as negative controls. Cells 48 hours after transfection were harvested and lysed in RIPA buffer with 1:100 Halt phosphatase cocktail inhibitor (Thermo Fisher 78420) and 1:50 Roche Complete protease inhibitor (11836145001) for 30 min on ice and cleared at 13,000 g for 15 min at 4 °C. Cell lysate was incubated with anti-FLAG (Sigma) at 4 °C with gentle agitation overnight, and then add 150 μl Protein A/G-coated magnetic beads incubation for 2 hours at 4 °C. The 40 μl 1 x loading buffer was added to the beads to extract proteins for SDS-PAGE.

### Flow cytometry analysis

Reporter cells were cultured for 5 days after transfection, trypsinized, washed with PBS, and resuspended in PBS for flow cytometry analysis (MACSQuant VYB). For each sample, 50,000 cells were analyzed. All data were analyzed by FlowJo 10.0 software.

### Western blotting

Cells were lysed 48 hours after transfection using cold radio immunoprecipitation assay (RIPA) buffer (Boston Bioproducts) supplemented with phosphatase inhibitor cocktail (Thermo Fisher Scientific) and protease inhibitor (Roche). Lysates were quantified using the bicinchoninic acid assay (BCA, Thermo Fisher Scientific). For each sample, 10 μg of protein were loaded onto a 15-well NuPAGE^™^ 4-12% Bis-Tris Protein Gels (Invitrogen), run at 100 V for 2.5 hours, then transferred to the nitrocellulose membrane (Thermo Fisher Scientific). The primary antibodies anti-ß-ACTIN (#4970, Cell Signaling Technology, dilution: 1:2000) or anti-Cas9 (A-9000-050, Epigentek Group, dilution: 1:2000) were used, followed by incubation with fluorophore-conjugated secondary antibodies. Bands were visualized using the Odyssey Imaging system (Li-Cor Biosciences).

### Deep sequencing and data analysis

Deep sequencing of genomic DNA was performed as previously described ^1^. For the first round of PCR, genomic sites of interest were amplified from 100 ng genomic DNA using Phusion Hot Start II PCR Master Mix with the primers containing Illumina forward and reverse adapters (listed in Supplementary Table 3). Briefly, 20 μL of PCR 1 reaction was performed with 0.5 μM of each forward and reverse primer, 1 μL genomic DNA extract, and 10 μL Phusion Flash PCR Master Mix (Thermo Fisher). PCR 1 reactions were carried out as follows: 98°C for 10 s, then 20 cycles of [98°C for 1 s, 55°C for 5 s, and 72°C for 6 s], followed by a final 72°C extension for 2 min. To add unique Illumina barcode, 30 μL of a given PCR 2 contained 1 μL unpurified PCR product, 0.5□μM of each unique forward and reverse Illumina barcoding primer pair, and 15 μL Phusion Flash PCR Master Mix. Primers are listed in Supplementary Table 3. PCR 2 reactions were carried out as follows: 98°C for 10 s, then 20 cycles of [98°C for 1 s, 60°C for 30 s, and 72°C for 6 s], followed by a final 72°C extension for 2 min. PCR 2 products were purified by electrophoresis with a 1% agarose gel using a QIAquick Gel Extraction Kit (Qiagen) and eluting with 30 μL water. DNA concentration was quantified by Qubit dsDNA HS Assay Kit (Thermo Fisher Scientific) or qPCR (KAPA Biosystems). The library was sequenced using an Illumina MiniSeq instrument according to the manufacturer’s protocols.

MiniSeq data analysis was done as previously reported ^1^. Sequencing reads were demultiplexed using bcl2fastq (Illumina). Alignment of amplicon reads to a reference sequence and prime editing efficiency calculations were performed using CRISPResso2 ^27^. To quantify the frequency of precise editing and indels, CRISPResso2 was run in standard mode with “discard_indel_reads” on. Precise editing efficiency was calculated as: (number of reads with precise edit) / (number of total reads), while indel efficiency was calculated as: 100% - precise editing efficiency - wildtype read rate.

### Readthrough assay

PE2, RT497 or empty vector were transfected with dual luciferase reporter in HEK293T cells. Cells were collected 48 h after transfection. readthrough assays were performed using the Dual-Glo Luciferase Assay System and following the manufacturer’s protocol (Promega, E2920).

### Animal studies

All animal experiments were approved by the Institutional Animal Care and Use Committee (IACUC) at UMass Medical School. All plasmids used for hydrodynamic tail-vein injection were prepared using EndoFree Plasmid Maxi kit (Qiagen). No randomization or blinding was used. For cancer model generation, eight-week-old FVB/NJ (Strain #001800) mice were injected with 2.5 mL saline containing 30 μg PE2 or 30 μg each of N-terminal cPE2 and C-terminal cPE2, 15 μg pegRNA, 15 μg nicking sgRNA, 5 μg pT3 EF1a-MYC (Addgene plasmid # 92046) and 1 μg CMV-SB10 (Addgene plasmid # 24551) via the tail vein in 5-7 s. 30 μg PE2, 15 μg non-target pegRNA, 5 μg pT3 EF1a-MYC and 1 μg CMV-SB10 were injected in the control group.

## Statistical analysis

GraphPad Prism 8 was used to analyze the data. All numerical values are presented as mean ± s.d.

## Data availability

The raw sequencing data have been deposited to the NCBI BioProject database. All raw data are available from the corresponding author upon request.

## Acknowledgements

We thank E. Sontheimer for helpful discussions and E. Haberlin for editing the manuscript. We thank Y. Liu in the Umass Morphology Core for support. This work was supported by grants from the National Institutes of Health (DP2HL137167, P01HL131471 and UG3HL147367), American Cancer Society (129056-RSG-16-093), the Lung Cancer Research Foundation, and the Cystic Fibrosis Foundation. P.L., and S.A.W. were supported in part by the National Institutes of Health (R01GM115911, and UG3TR002668) and the Rett Syndrome Research Trust.

## Authors’ contributions

W.X., C.Z., and S.A.W. designed the research; C.Z. performed experiments, analyzed the data and wrote the manuscript. S.L., B.L., and P.L. generated cell report lines and did pegRNA clone. All authors reviewed, edited, and approved the manuscript.

## Supplementary data accompanies this paper

