## Supplementary material for "Development of a flexible split prime editor using truncated reverse transcriptase": Supplementary Data.docx

Supplementary Fig.1

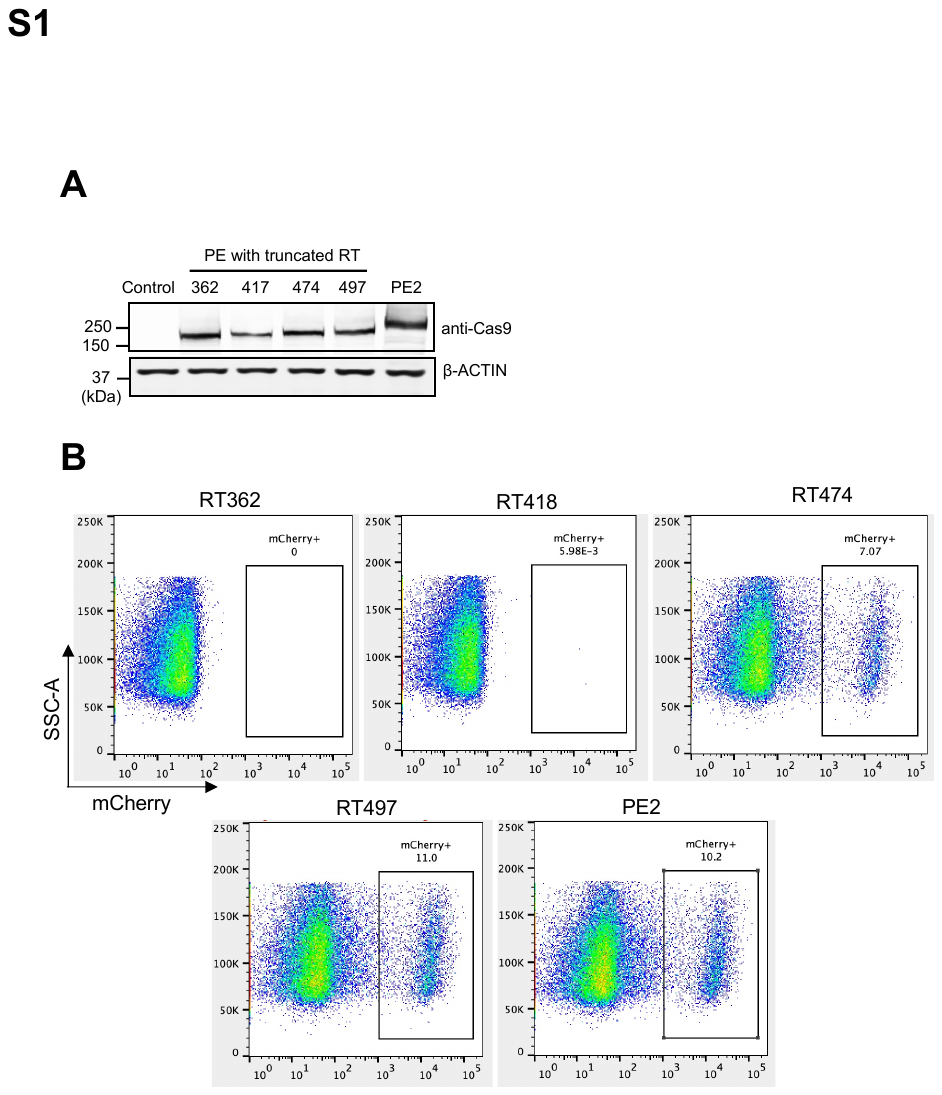

Supplementary Fig. 1: Flow cytometry analysis of editing efficiency of full-length PE2 and RT variants.

a Western blot of full-length PE2 and compact PE2 variants. b Representative data of flow cytometry analysis of mCherry+ cells. The image data was analyzed by FlowJo 10.0 software. HEK293T cells were initially gated using FSC-A/SSC-A, then sorted for single cell using FSC-A/FSC-H. mCherry-positive cells were gated by SSC-A/Y2-A.

Supplementary Fig.2

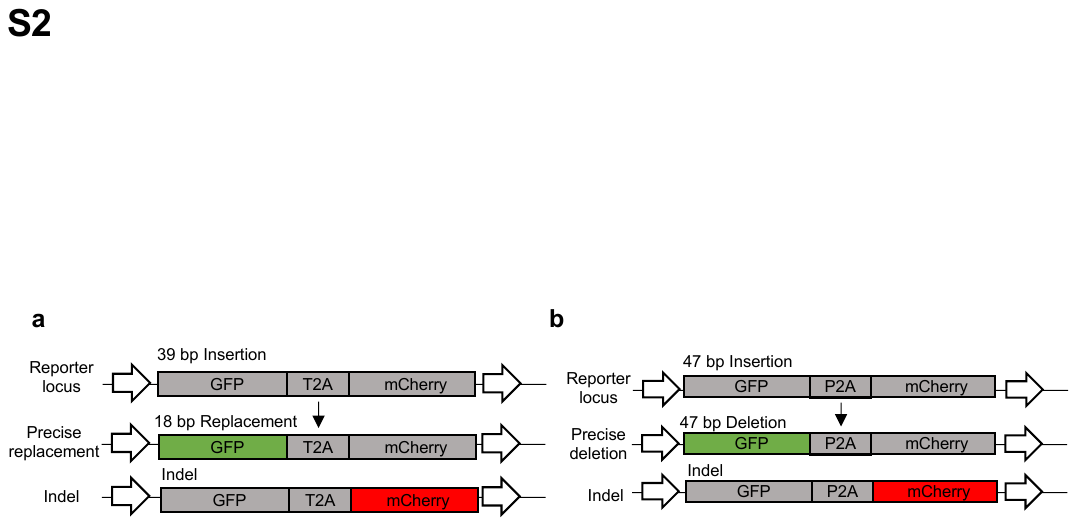

Supplementary Fig. 2 Editing frequencies of RT variants in TLR-MCV1 and TLR reporter lines.

**a** Traffic light reporter multi-cas variant 1 (TLR-MCV1) cells containing a GFP with a 39-bp insertion, P2A, and out-of-frame mCherry. **b** TLR system containing a GFP with 47-bp insertion.

Supplementary Fig.3

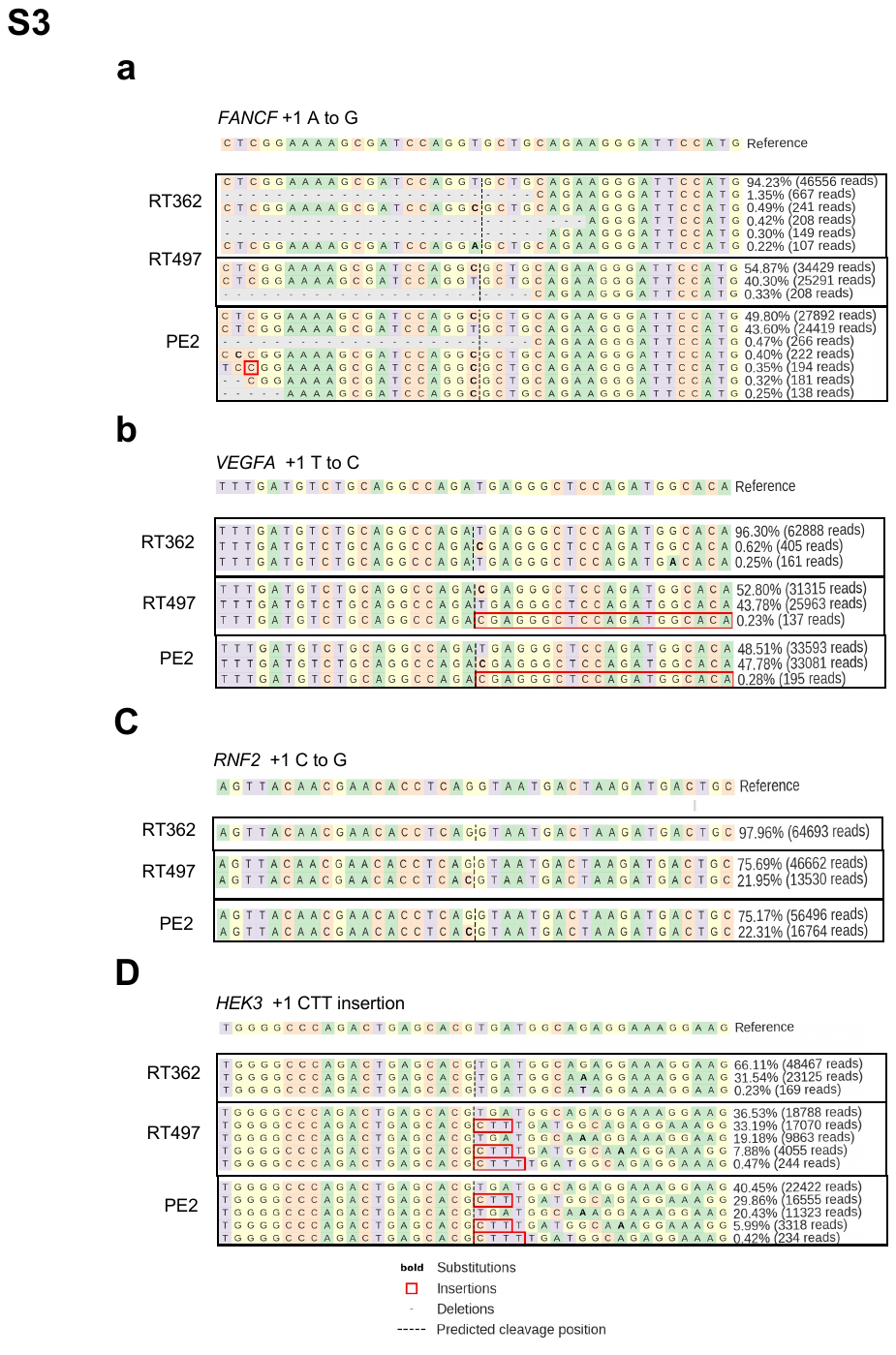

Supplementary Fig. 3: Deep sequencing of endogenous loci.

**a-d** Allele frequencies and corresponding Illumina sequencing read counts are shown for each allele. All alleles observed with frequency ≥ 0.2% are shown.

Supplementary Fig.4

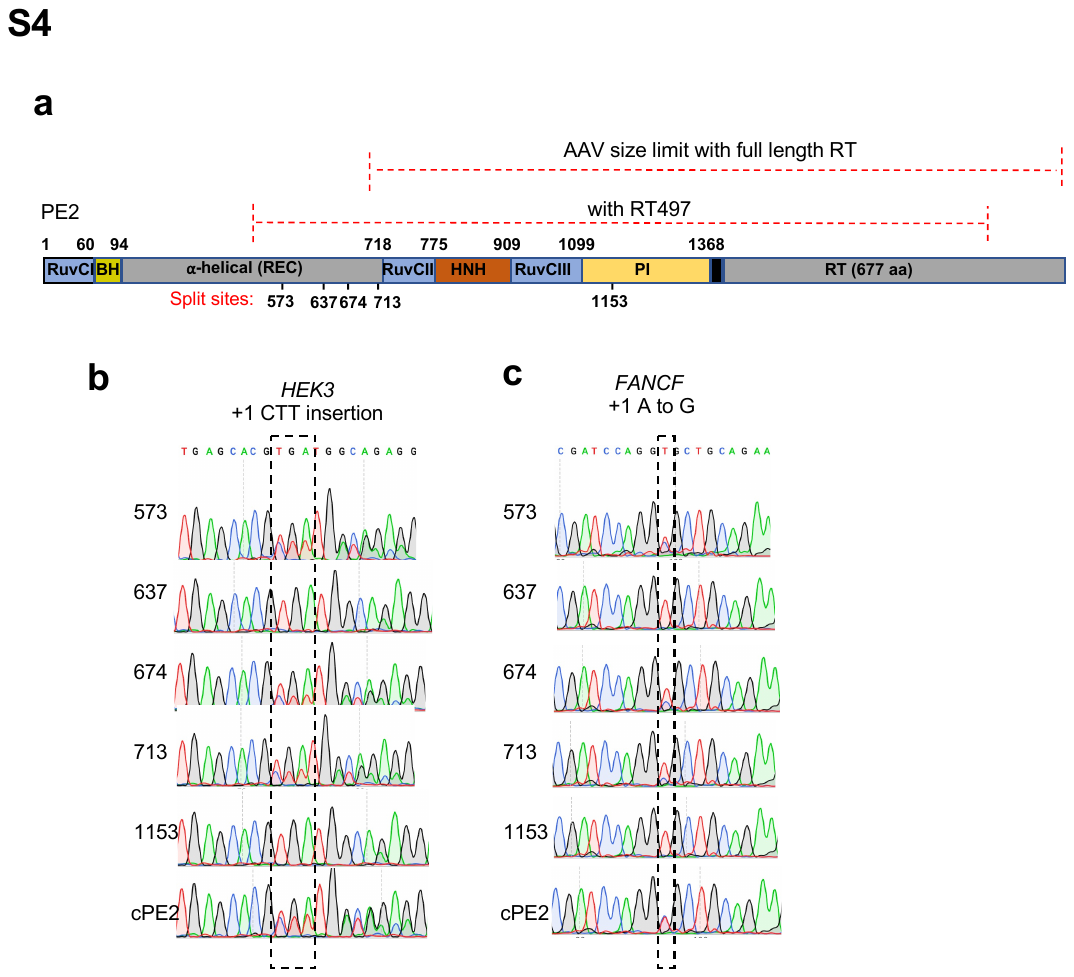

Supplementary Fig. 4: Sanger sequencing of endogenous loci of split-cPE2 and cPE2.

**a** Schematic representation of PE2 and cPE2 split sites. AAV size limit is ~4.7 kb without ITR sequence. **b-c** Sanger sequencing showed +1 CTT insertion at *HEK3* locus **b** and +1 A to G transversion at *FANCF* locus. **c** Split-cPE2-573 supports robust editing efficiency at *HEK3* and *FANCF* loci.

Supplementary Fig.5

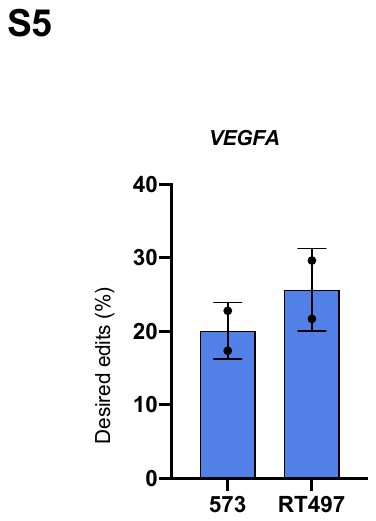

Supplementary Fig. 5: Editing frequencies of split-cPE2 at VEGFA.

**Supplementary Table 1**. Sequences of pegRNAs and sgRNAs used in this study. All sequences are shown in 5' to 3' orientation.

**Supplementary Table 2**. Sequences of primers used for cloning.

**Supplementary Table 3**. Sequences of primers used for genomic DNA amplification and high throughput sequencing.

**Supplementary Sequences:** Sequence of backbone plasmid used for prime editing

**Supplementary Table 1**. Sequences of pegRNAs and sgRNAs used in this study. All sequences are shown in 5' to 3' orientation.

sgRNA scaffold GTTTTAGAGCTAGAAATAGCAAGTTAAAATAAGGCTAGTCCGTTATCAACTTGAAAAAGTGGGACCGAGTCGGTCC

| **pegRNA** | **spacer sequence (5'-3')** | **3' extension** | **PBS (nt)** | **RT (nt)** | **Figure** |
| --- | --- | --- | --- | --- | --- |
| mCherry A to G | CACCTTCAGCTTGGCGGTCT | TACGAGGGCACTCAAACCGCCAAGCTGAAG | 14 | 16 | Figure 1C |
| GFP-insertion | AAGTTCAGCGTGTCCGGCTT | GTCAGCTTGCCGTAGGTGGCATCGCCCTCGCCTTCG | 13 | 36 | Figure 1D |
| GFP-deletion | GCGGAGAGGGCACCCCCGA | GTTGGTCATGCGACCCTGCTCGGGGGTGCCCTCTCC | 14 | 22 | Figure 1E |
| FANCF +1 A to G | GGAATCCCTTCTGCAGCACC | GGAAAAGCGATCCAGGCGCTGCAGAAGGGAT | 14 | 17 | Figure 2A, 3B |
| VEGFA +1 T to C | GATGTCTGCAGGCCAGATGA | AATGTGCCATCTGGAGCCCTCGTCTGGCCTGCAGA | 13 | 22 | Figure 2A |
| RNF2 +1 C to G | GTCATCTTAGTCATTACCTG | AACGAACACCTCACGTAATGACTAAGATG | 15 | 14 | Figure 2A |
| HEK3 +1 CTT insertion | GGCCCAGACTGAGCACGTGA | TCTGCCATCAAAGCGTGCTCAGTCTG | 13 | 13 | Figure 2A, 3B |
| HEK3 +1 T to A | GGCCCAGACTGAGCACGTGA | TGGAGGAAGCAGGGCTTCCTTTCCTCTGCCATCTCGTGCTCAGTCTG | 13 | 34 | Figure 2B |
| HEK3 +12 G to C | GGCCCAGACTGAGCACGTGA | TGGAGGAAGCAGGGCTTCCTTTGCTCTGCCATCACGTGCTCAGTCTG | 13 | 34 | Figure 2B |
| HEK3 +30 C to G | GGCCCAGACTGAGCACGTGA | TGGACGAAGCAGGGCTTCCTTTCCTCTGCCATCACGTGCTCAGTCTG | 13 | 34 | Figure 2B |
| Ctnnb1 C to T | AGGGTTGCCCTTGCCACTCA | GCTCCTTTCCTGAGTGGCAAGGGCAA | 13 | 13 | Figure 3C |

| **Nicking sgRNA** | **spacer sequence (5'-3')** | **Figure** |
| --- | --- | --- |
| mCherry A to G | GCTGTCCCCTCAGTTCATGTA | Figure 1C |
| GFP-insertion | GTAGGTCAGGGTGGTCACGA | Figure 1D |
| GFP-deletion | GAGAAGCCGTAGCCCATCACG | Figure 1E |
| FANCF +1 A to G | GGGGTCCCAGGTGCTGACGT | Figure 2A, 3B |
| VEGFA +1 T to C | GATGTACAGAGAGCCCAGGGC | Figure 2A |
| RNF2 +1 C to G | GTCATCTTAGTCATTACCTG | Figure 2A |
| HEK3 +1 CTT insertion | GGCCCAGACTGAGCACGTGA | Figure 2A, 3B |
| HEK3 +1 T to A | GGCCCAGACTGAGCACGTGA | Figure 2B |
| HEK3 +12 G to C | GGCCCAGACTGAGCACGTGA | Figure 2B |
| HEK3 +30 C to G | GGCCCAGACTGAGCACGTGA | Figure 2B |
| Ctnnb1 C to T | GAAAAGCTGCTGTCAGCCAC | Figure 3C |

**Supplementary Table 2**. Sequences of primers used for cloning.

|  | **F (5'-3')** | **R (5'-3')** |
| --- | --- | --- |
| RT362 | CAGATTTGTCTGGCGGCTCAAAAAGAACC | CCGCCAGACAAATCTGGCAACCCCAGGG |
| RT418 | TTGCCGTATCTGGCGGCTCAAAAAGAACC | CCGCCAGATACGGCAATGGCTGCTACC |
| RT474 | TCGGACCGTCTGGCGGCTCAAAAAGAACC | CCGCCAGACGGTCCGAACTGGACCCGG |
| RT497 | GCCTTGATTCTGGCGGCTCAAAAAGAACC | CCGCCAGAATCAAGGCAGTTGTGTTGCA |
| N-terminal Split-cPE2-573 | TCGAGTCACCAAAGAAGAAGCGGAAAGTCGACAAGAAGTACAGCATCGGC | TGTCAGGATCTCTGTCTCGTAGGACAGGCACTCGATTTTCTTGAAGTAGTCCTC |
|  | TGCCTGTCCTACGAGACAGA | GCCGTCGGCGGTTCTTTTTGAGCCGCCAGAATTAGGCAGGTTATCCACTC |
|  | TCTGGCGGCTCAAAAAGAAC | GACTTTCCGCTTCTTCTTTGG |
| C-terminal Split-cPE2-573 | TTCGAGTCACCAAAGAAGAAGCGGAAAGTCATGATCAAGATTGCTACACG | CACGCCGGAGATTTCCACGGAGTCGAAGCAATTGCTGGCGATAAAGCCA |
|  | TGCTTCGACTCCGTGGAAAT | GCCGTCGGCGGTTCTTTTTGAGCCGCCAGAATCAAGGCAGTTGTGTTGCA |
|  | TCTGGCGGCTCAAAAAGAAC | GACTTTCCGCTTCTTCTTTGG |
| N-terminal Split-cPE2-637 | TCGAGTCACCAAAGAAGAAGCGGAAAGTCGACAAGAAGTACAGCATCGGC | TGTCAGGATCTCTGTCTCGTAGGACAGGCATTTCAGCCGTTCCTCGATCA |
|  | TGCCTGTCCTACGAGACAGA | GCCGTCGGCGGTTCTTTTTGAGCCGCCAGAATTAGGCAGGTTATCCACTC |
|  | TCTGGCGGCTCAAAAAGAAC | GACTTTCCGCTTCTTCTTTGG |
| C-terminal Split-cPE2-637 | TTCGAGTCACCAAAGAAGAAGCGGAAAGTCATGATCAAGATTGCTACACG | ATTGCTGGCGATAAAGCCAT |
|  | ACCTATGCCCACCTGTTCGA | GCCGTCGGCGGTTCTTTTTGAGCCGCCAGAATCAAGGCAGTTGTGTTGCA |
|  | TCTGGCGGCTCAAAAAGAAC | GACTTTCCGCTTCTTCTTTGG |
| N-terminal Split-cPE2-674 | TCGAGTCACCAAAGAAGAAGCGGAAAGTCGACAAGAAGTACAGCATCGGC | TGTCAGGATCTCTGTCTCGTAGGACAGGCACTGCTTGTCCCGGATGCCG |
|  | TGCCTGTCCTACGAGACAGA | GCCGTCGGCGGTTCTTTTTGAGCCGCCAGAATTAGGCAGGTTATCCACTC |
|  | TCTGGCGGCTCAAAAAGAAC | GACTTTCCGCTTCTTCTTTGG |
| C-terminal Split-cPE2-674 | TTCGAGTCACCAAAGAAGAAGCGGAAAGTCATGATCAAGATTGCTACACG | ATTGCTGGCGATAAAGCCAT |
|  | GCCCTGAAGAATGGCTTTATCGCCAGCAATTCCGGCAAGACAATCCTGGA | GCCGTCGGCGGTTCTTTTTGAGCCGCCAGAATCAAGGCAGTTGTGTTGCA |
|  | TCTGGCGGCTCAAAAAGAAC | GACTTTCCGCTTCTTCTTTGG |
| N-terminal Split-cPE2-713 | TCGAGTCACCAAAGAAGAAGCGGAAAGTCGACAAGAAGTACAGCATCGGC | TGTCAGGATCTCTGTCTCGTAGGACAGGCAcacctgggctttctggatgt |
|  | TGCCTGTCCTACGAGACAGA | GCCGTCGGCGGTTCTTTTTGAGCCGCCAGAATTAGGCAGGTTATCCACTC |
|  | TCTGGCGGCTCAAAAAGAAC | GACTTTCCGCTTCTTCTTTGG |
| C-terminal Split-cPE2-713 | TTCGAGTCACCAAAGAAGAAGCGGAAAGTCATGATCAAGATTGCTACACG | ATTGCTGGCGATAAAGCCAT |
|  | GCCCTGAAGAATGGCTTTATCGCCAGCAATtccggccagggcgatagcc | GCCGTCGGCGGTTCTTTTTGAGCCGCCAGAATCAAGGCAGTTGTGTTGCA |
|  | TCTGGCGGCTCAAAAAGAAC | GACTTTCCGCTTCTTCTTTGG |
| N-terminal Split-cPE2-1153 | TCGAGTCACCAAAGAAGAAGCGGAAAGTCGACAAGAAGTACAGCATCGGC | TGTCAGGATCTCTGTCTCGTAGGACAGGCACTTGCCCTTTTCCACTTTGG |
|  | TGCCTGTCCTACGAGACAGA | GCCGTCGGCGGTTCTTTTTGAGCCGCCAGAATTAGGCAGGTTATCCACTC |
|  | TCTGGCGGCTCAAAAAGAAC | GACTTTCCGCTTCTTCTTTGG |
| C-terminal Split-cPE2-1153 | TTCGAGTCACCAAAGAAGAAGCGGAAAGTCATGATCAAGATTGCTACACG | ATTGCTGGCGATAAAGCCAT |
|  | CCCTGAAGAATGGCTTTATCGCCAGCAATTCCAAGAAACTGAAGAGTGTG | GCCGTCGGCGGTTCTTTTTGAGCCGCCAGAATCAAGGCAGTTGTGTTGCA |
|  | TCTGGCGGCTCAAAAAGAAC | GACTTTCCGCTTCTTCTTTGG |

**Supplementary Table 3**. Sequences of primers used for genomic DNA amplification and high throughput sequencing.

| **Gene** | **F (5'-3')** | **R (5'-3')** | **Figure** |
| --- | --- | --- | --- |
| FANCF | CTACACGACGCTCTTCCGATCTGATGGATGTGGCGCAGGTAG | AGACGTGTGCTCTTCCGATCTAGGCGTATCATTTCGCGGAT | Figure 2A, 3B |
| VEGFA | CTACACGACGCTCTTCCGATCTAAGCATCCCTGGACACTTCC | AGACGTGTGCTCTTCCGATCTTGGACCCCCTATTTCTGACCT | Figure 2A |
| RNF2 | CTACACGACGCTCTTCCGATCTACCATAGCACTTCCCTTCCA | AGACGTGTGCTCTTCCGATCTTATCCCAGTTTACACGTCTC | Figure 2A |
| HEK3 | CTACACGACGCTCTTCCGATCTTGCTGCAAGTA  AGCATGCATTTG | AGACGTGTGCTCTTCCGATCTCTTCCAGCCCAGCCAAACTT | Figure 2A, 2B, 3B |

**Supplementary Sequences**  Sequence of backbone plasmid used for prime editing

Sequence of reverse transcriptase: Finger-Palm domain (1-275 aa) + Thumb domain (276-361 aa) + Connection domain (362 - 496 aa) + RNase H domain (497-671 aa)

TLNIEDEYRLHETSKEPDVSLGSTWLSDFPQAWAETGGMGLAVRQAPLIIPLKATSTPVSIKQYPMSQEARLGIKPHIQRLLDQGILVPCQSPWNTPLLPVKKPGTNDYRPVQDLREVNKRVEDIHPTVPNPYNLLSGLPPSHQWYTVLDLKDAFFCLRLHPTSQPLFAFEWRDPEMGISGQLTWTRLPQGFKNSPTLFNEALHRDLADFRIQHPDLILLQYVDDLLLAATSELDCQQGTRALLQTLGNLGYRASAKKAQICQKQVKYLGYLLKEGQRWLTEARKETVMGQPTPKTPRQLREFLGTAGFCRLWIPGFAEMAAPLYPLTKPGTLFNWGPDQQKAYQEIKQALLTAPALGLPDLTKPFELFVDEKQGYAKGVLTQKLGPWRRPVAYLSKKLDPVAAGWPPCLRMVAAIAVLTKDAGKLTMGQPLVILAPHAVEALVKQPPDRWLSNARMTHYQALLLDTDRVQFGPVVALNPATLLPLPEEGLQHNCLDILAEAHGTRPDLTDQPLPDADHTWYTDGSSLLQEGQRKAGAAVTTETEVIWAKALPAGTSAQRAELIALTQALKMAEGKKLNVYTDSRYAFATAHIHGEIYRRRGWLTSEGKEIKNKDEILALLKALFLPKRLSIIHCPGHQKGHSAEARGNRMADQAARKAAITETPDTSTLL

Sequence of RT497: cmv promoter + N-terminal NLS + Cas9 H840A + Flexible linker + MoMLV reverse transcriptase + C-terminal NLS + Plasmid backbone

gacattgattattgactagttattaatagtaatcaattacggggtcattagttcatagcccatatatggagttccgcgttacataacttacggtaaatggcccgcctggctgaccgcccaacgacccccgcccattgacgtcaataatgacgtatgttcccatagtaacgccaatagggactttccattgacgtcaatgggtggagtatttacggtaaactgcccacttggcagtacatcaagtgtatcatatgccaagtacgccccctattgacgtcaatgacggtaaatggcccgcctggcattatgcccagtacatgaccttatgggactttcctacttggcagtacatctacgtattagtcatcgctattaccatggtgatgcggttttggcagtacatcaatgggcgtggatagcggtttgactcacggggatttccaagtctccaccccattgacgtcaatgggagtttgttttggcaccaaaatcaacgggactttccaaaatgtcgtaacaactccgccccattgacgcaaatgggcggtaggcgtgtacggtgggaggtctatataagcagagctggtttagtgaaccgtcagatccgctagagatccgcggccgctaatacgactcactatagggagagccgccaccatgaaacggacagccgacggaagcgagttcgagtcaccaaagaagaagcggaaagtcgacaagaagtacagcatcggcctggacatcggcaccaactctgtgggctgggccgtgatcaccgacgagtacaaggtgcccagcaagaaattcaaggtgctgggcaacaccgaccggcacagcatcaagaagaacctgatcggagccctgctgttcgacagcggcgaaacagccgaggccacccggctgaagagaaccgccagaagaagatacaccagacggaagaaccggatctgctatctgcaagagatcttcagcaacgagatggccaaggtggacgacagcttcttccacagactggaagagtccttcctggtggaagaggataagaagcacgagcggcaccccatcttcggcaacatcgtggacgaggtggcctaccacgagaagtaccccaccatctaccacctgagaaagaaactggtggacagcaccgacaaggccgacctgcggctgatctatctggccctggcccacatgatcaagttccggggccacttcctgatcgagggcgacctgaaccccgacaacagcgacgtggacaagctgttcatccagctggtgcagacctacaaccagctgttcgaggaaaaccccatcaacgccagcggcgtggacgccaaggccatcctgtctgccagactgagcaagagcagacggctggaaaatctgatcgcccagctgcccggcgagaagaagaatggcctgttcggaaacctgattgccctgagcctgggcctgacccccaacttcaagagcaacttcgacctggccgaggatgccaaactgcagctgagcaaggacacctacgacgacgacctggacaacctgctggcccagatcggcgaccagtacgccgacctgtttctggccgccaagaacctgtccgacgccatcctgctgagcgacatcctgagagtgaacaccgagatcaccaaggcccccctgagcgcctctatgatcaagagatacgacgagcaccaccaggacctgaccctgctgaaagctctcgtgcggcagcagctgcctgagaagtacaaagagattttcttcgaccagagcaagaacggctacgccggctacattgacggcggagccagccaggaagagttctacaagttcatcaagcccatcctggaaaagatggacggcaccgaggaactgctcgtgaagctgaacagagaggacctgctgcggaagcagcggaccttcgacaacggcagcatcccccaccagatccacctgggagagctgcacgccattctgcggcggcaggaagatttttacccattcctgaaggacaaccgggaaaagatcgagaagatcctgaccttccgcatcccctactacgtgggccctctggccaggggaaacagcagattcgcctggatgaccagaaagagcgaggaaaccatcaccccctggaacttcgaggaagtggtggacaagggcgcttccgcccagagcttcatcgagcggatgaccaacttcgataagaacctgcccaacgagaaggtgctgcccaagcacagcctgctgtacgagtacttcaccgtgtataacgagctgaccaaagtgaaatacgtgaccgagggaatgagaaagcccgccttcctgagcggcgagcagaaaaaggccatcgtggacctgctgttcaagaccaaccggaaagtgaccgtgaagcagctgaaagaggactacttcaagaaaatcgagtgcttcgactccgtggaaatctccggcgtggaagatcggttcaacgcctccctgggcacataccacgatctgctgaaaattatcaaggacaaggacttcctggacaatgaggaaaacgaggacattctggaagatatcgtgctgaccctgacactgtttgaggacagagagatgatcgaggaacggctgaaaacctatgcccacctgttcgacgacaaagtgatgaagcagctgaagcggcggagatacaccggctggggcaggctgagccggaagctgatcaacggcatccgggacaagcagtccggcaagacaatcctggatttcctgaagtccgacggcttcgccaacagaaacttcatgcagctgatccacgacgacagcctgacctttaaagaggacatccagaaagcccaggtgtccggccagggcgatagcctgcacgagcacattgccaatctggccggcagccccgccattaagaagggcatcctgcagacagtgaaggtggtggacgagctcgtgaaagtgatgggccggcacaagcccgagaacatcgtgatcgaaatggccagagagaaccagaccacccagaagggacagaagaacagccgcgagagaatgaagcggatcgaagagggcatcaaagagctgggcagccagatcctgaaagaacaccccgtggaaaacacccagctgcagaacgagaagctgtacctgtactacctgcagaatgggcgggatatgtacgtggaccaggaactggacatcaaccggctgtccgactacgatgtggacgctatcgtgcctcagagctttctgaaggacgactccatcgacaacaaggtgctgaccagaagcgacaagaaccggggcaagagcgacaacgtgccctccgaagaggtcgtgaagaagatgaagaactactggcggcagctgctgaacgccaagctgattacccagagaaagttcgacaatctgaccaaggccgagagaggcggcctgagcgaactggataaggccggcttcatcaagagacagctggtggaaacccggcagatcacaaagcacgtggcacagatcctggactcccggatgaacactaagtacgacgagaatgacaagctgatccgggaagtgaaagtgatcaccctgaagtccaagctggtgtccgatttccggaaggatttccagttttacaaagtgcgcgagatcaacaactaccaccacgcccacgacgcctacctgaacgccgtcgtgggaaccgccctgatcaaaaagtaccctaagctggaaagcgagttcgtgtacggcgactacaaggtgtacgacgtgcggaagatgatcgccaagagcgagcaggaaatcggcaaggctaccgccaagtacttcttctacagcaacatcatgaactttttcaagaccgagattaccctggccaacggcgagatccggaagcggcctctgatcgagacaaacggcgaaaccggggagatcgtgtgggataagggccgggattttgccaccgtgcggaaagtgctgagcatgccccaagtgaatatcgtgaaaaagaccgaggtgcagacaggcggcttcagcaaagagtctatcctgcccaagaggaacagcgataagctgatcgccagaaagaaggactgggaccctaagaagtacggcggcttcgacagccccaccgtggcctattctgtgctggtggtggccaaagtggaaaagggcaagtccaagaaactgaagagtgtgaaagagctgctggggatcaccatcatggaaagaagcagcttcgagaagaatcccatcgactttctggaagccaagggctacaaagaagtgaaaaaggacctgatcatcaagctgcctaagtactccctgttcgagctggaaaacggccggaagagaatgctggcctctgccggcgaactgcagaagggaaacgaactggccctgccctccaaatatgtgaacttcctgtacctggccagccactatgagaagctgaagggctcccccgaggataatgagcagaaacagctgtttgtggaacagcacaagcactacctggacgagatcatcgagcagatcagcgagttctccaagagagtgatcctggccgacgctaatctggacaaagtgctgtccgcctacaacaagcaccgggataagcccatcagagagcaggccgagaatatcatccacctgtttaccctgaccaatctgggagcccctgccgccttcaagtactttgacaccaccatcgaccggaagaggtacaccagcaccaaagaggtgctggacgccaccctgatccaccagagcatcaccggcctgtacgagacacggatcgacctgtctcagctgggaggtgactctggaggatctagcggaggatcctctggcagcgagacaccaggaacaagcgagtcagcaacaccagagagcagtggcggcagcagcggcggcagcagcaccctaaatatagaagatgagtatcggctacatgagacctcaaaagagccagatgtttctctagggtccacatggctgtctgattttcctcaggcctgggcggaaaccgggggcatgggactggcagttcgccaagctcctctgatcatacctctgaaagcaacctctacccccgtgtccataaaacaataccccatgtcacaagaagccagactggggatcaagccccacatacagagactgttggaccagggaatactggtaccctgccagtccccctggaacacgcccctgctacccgttaagaaaccagggactaatgattataggcctgtccaggatctgagagaagtcaacaagcgggtggaagacatccaccccaccgtgcccaacccttacaacctcttgagcgggctcccaccgtcccaccagtggtacactgtgcttgatttaaaggatgcctttttctgcctgagactccaccccaccagtcagcctctcttcgcctttgagtggagagatccagagatgggaatctcaggacaattgacctggaccagactcccacagggtttcaaaaacagtcccaccctgtttaatgaggcactgcacagagacctagcagacttccggatccagcacccagacttgatcctgctacagtacgtggatgacttactgctggccgccacttctgagctagactgccaacaaggtactcgggccctgttacaaaccctagggaacctcgggtatcgggcctcggccaagaaagcccaaatttgccagaaacaggtcaagtatctggggtatcttctaaaagagggtcagagatggctgactgaggccagaaaagagactgtgatggggcagcctactccgaagacccctcgacaactaagggagttcctagggaaggcaggcttctgtcgcctcttcatccctgggtttgcagaaatggcagcccccctgtaccctctcaccaaaccggggactctgtttaattggggcccagaccaacaaaaggcctatcaagaaatcaagcaagctcttctaactgccccagccctggggttgccagatttgactaagccctttgaactctttgtcgacgagaagcagggctacgccaaaggtgtcctaacgcaaaaactgggaccttggcgtcggccggtggcctacctgtccaaaaagctagacccagtagcagctgggtggcccccttgcctacggatggtagcagccattgccgtactgacaaaggatgcaggcaagctaaccatgggacagccactagtcattctggccccccatgcagtagaggcactagtcaaacaaccccccgaccgctggctttccaacgcccggatgactcactatcaggccttgcttttggacacggaccgggtccagttcggaccggtggtagccctgaacccggctacgctgctcccactgcctgaggaagggctgcaacacaactgccttgattctggcggctcaaaaagaaccgccgacggcagcgaattcgagcccaagaagaagaggaaagtctaaccggtcatcatcaccatcaccattgagtttaaacccgctgatcagcctcgactgtgccttctagttgccagccatctgttgtttgcccctcccccgtgccttccttgaccctggaaggtgccactcccactgtcctttcctaataaaatgagaaaattgcatcgcattgtctgagtaggtgtcattctattctggggggtggggtggggcaggacagcaagggggaggattgggaagacaatagcaggcatgctggggatgcggtgggctctatggcttctgaggcggaaagaaccagctggggctcgataccgtcgacctctagctagagcttggcgtaatcatggtcatagctgtttcctgtgtgaaattgttatccgctcacaattccacacaacatacgagccggaagcataaagtgtaaagcctagggtgcctaatgagtgagctaactcacattaattgcgttgcgctcactgcccgctttccagtcgggaaacctgtcgtgccagctgcattaatgaatcggccaacgcgcggggagaggcggtttgcgtattgggcgctcttccgcttcctcgctcactgactcgctgcgctcggtcgttcggctgcggcgagcggtatcagctcactcaaaggcggtaatacggttatccacagaatcaggggataacgcaggaaagaacatgtgagcaaaaggccagcaaaaggccaggaaccgtaaaaaggccgcgttgctggcgtttttccataggctccgcccccctgacgagcatcacaaaaatcgacgctcaagtcagaggtggcgaaacccgacaggactataaagataccaggcgtttccccctggaagctccctcgtgcgctctcctgttccgaccctgccgcttaccggatacctgtccgcctttctcccttcgggaagcgtggcgctttctcatagctcacgctgtaggtatctcagttcggtgtaggtcgttcgctccaagctgggctgtgtgcacgaaccccccgttcagcccgaccgctgcgccttatccggtaactatcgtcttgagtccaacccggtaagacacgacttatcgccactggcagcagccactggtaacaggattagcagagcgaggtatgtaggcggtgctacagagttcttgaagtggtggcctaactacggctacactagaagaacagtatttggtatctgcgctctgctgaagccagttaccttcggaaaaagagttggtagctcttgatccggcaaacaaaccaccgctggtagcggtggtttttttgtttgcaagcagcagattacgcgcagaaaaaaaggatctcaagaagatcctttgatcttttctacggggtctgacactcagtggaacgaaaactcacgttaagggattttggtcatgagattatcaaaaaggatcttcacctagatccttttaaattaaaaatgaagttttaaatcaatctaaagtatatatgagtaaacttggtctgacagttaccaatgcttaatcagtgaggcacctatctcagcgatctgtctatttcgttcatccatagttgcctgactccccgtcgtgtagataactacgatacgggagggcttaccatctggccccagtgctgcaatgataccgcgagacccacgctcaccggctccagatttatcagcaataaaccagccagccggaagggccgagcgcagaagtggtcctgcaactttatccgcctccatccagtctattaattgttgccgggaagctagagtaagtagttcgccagttaatagtttgcgcaacgttgttgccattgctacaggcatcgtggtgtcacgctcgtcgtttggtatggcttcattcagctccggttcccaacgatcaaggcgagttacatgatcccccatgttgtgcaaaaaagcggttagctccttcggtcctccgatcgttgtcagaagtaagttggccgcagtgttatcactcatggttatggcagcactgcataattctcttactgtcatgccatccgtaagatgcttttctgtgactggtgagtactcaaccaagtcattctgagaatagtgtatgcggcgaccgagttgctcttgcccggcgtcaatacgggataataccgcgccacatagcagaactttaaaagtgctcatcattggaaaacgttcttcggggcgaaaactctcaaggatcttaccgctgttgagatccagttcgatgtaacccactcgtgcacccaactgatcttcagcatcttttactttcaccagcgtttctgggtgagcaaaaacaggaaggcaaaatgccgcaaaaaagggaataagggcgacacggaaatgttgaatactcatactcttcctttttcaatattattgaagcatttatcagggttattgtctcatgagcggatacatatttgaatgtatttagaaaaataaacaaataggggttccgcgcacatttccccgaaaagtgccacctgacgtcgacggatcgggagatcgatctcccgatcccctagggtctactctcagtacaatctgctctgatgccgcatagttaagccagtatctgctccctgcttgtgtgttggaggtcgctgagtagtgcgcgagcaaaatttaagctacaacaaggcaaggcttgaccgacaattgcatgaagaatctgcttagggttaggcgttttgcgctgcttcgcgatgtacgggccagatatacgcgtt

Backbone of pegRNA and Nicking sgRNA: U6 promoter + spCas9-sgRNA scaffold + Plasmid backbone

GTGGCACTTTTCGGGGAAATGTGCGCGGAACCCCTATTTGTTTATTTTTCTAAATACATTCAAATATGTATCCGCTCATGAGACAATAACCCTGATAAATGCTTCAATAATATTGAAAAAGGAAGAGTATGAGTATTCAACATTTCCGTGTCGCCCTTATTCCCTTTTTTGCGGCATTTTGCCTTCCTGTTTTTGCTCACCCAGAAACGCTGGTGAAAGTAAAAGATGCTGAAGATCAGTTGGGTGCACGAGTGGGTTACATCGAACTGGATCTCAACAGCGGTAAGATCCTTGAGAGTTTTCGCCCCGAAGAACGTTTTCCAATGATGAGCACTTTTAAAGTTCTGCTATGTGGCGCGGTATTATCCCGTATTGACGCCGGGCAAGAGCAACTCGGTCGCCGCATACACTATTCTCAGAATGACTTGGTTGAGTACTCACCAGTCACAGAAAAGCATCTTACGGATGGCATGACAGTAAGAGAATTATGCAGTGCTGCCATAACCATGAGTGATAACACTGCGGCCAACTTACTTCTGACAACGATCGGAGGACCGAAGGAGCTAACCGCTTTTTTGCACAACATGGGGGATCATGTAACTCGCCTTGATCGTTGGGAACCGGAGCTGAATGAAGCCATACCAAACGACGAGCGTGACACCACGATGCCTGTAGCAATGGCAACAACGTTGCGCAAACTATTAACTGGCGAACTACTTACTCTAGCTTCCCGGCAACAATTAATAGACTGGATGGAGGCGGATAAAGTTGCAGGACCACTTCTGCGCTCGGCCCTTCCGGCTGGCTGGTTTATTGCTGATAAATCTGGAGCCGGTGAGCGTGGGTCTCGCGGTATCATTGCAGCACTGGGGCCAGATGGTAAGCCCTCCCGTATCGTAGTTATCTACACGACGGGGAGTCAGGCAACTATGGATGAACGAAATAGACAGATCGCTGAGATAGGTGCCTCACTGATTAAGCATTGGTAACTGTCAGACCAAGTTTACTCATATATACTTTAGATTGATTTAAAACTTCATTTTTAATTTAAAAGGATCTAGGTGAAGATCCTTTTTGATAATCTCATGACCAAAATCCCTTAACGTGAGTTTTCGTTCCACTGAGCGTCAGACCCCGTAGAAAAGATCAAAGGATCTTCTTGAGATCCTTTTTTTCTGCGCGTAATCTGCTGCTTGCAAACAAAAAAACCACCGCTACCAGCGGTGGTTTGTTTGCCGGATCAAGAGCTACCAACTCTTTTTCCGAAGGTAACTGGCTTCAGCAGAGCGCAGATACCAAATACTGTCCTTCTAGTGTAGCCGTAGTTAGGCCACCACTTCAAGAACTCTGTAGCACCGCCTACATACCTCGCTCTGCTAATCCTGTTACCAGTGGCTGCTGCCAGTGGCGATAAGTCGTGTCTTACCGGGTTGGACTCAAGACGATAGTTACCGGATAAGGCGCAGCGGTCGGGCTGAACGGGGGGTTCGTGCACACAGCCCAGCTTGGAGCGAACGACCTACACCGAACTGAGATACCTACAGCGTGAGCTATGAGAAAGCGCCACGCTTCCCGAAGGGAGAAAGGCGGACAGGTATCCGGTAAGCGGCAGGGTCGGAACAGGAGAGCGCACGAGGGAGCTTCCAGGGGGAAACGCCTGGTATCTTTATAGTCCTGTCGGGTTTCGCCACCTCTGACTTGAGCGTCGATTTTTGTGATGCTCGTCAGGGGGGCGGAGCCTATGGAAAAACGCCAGCAACGCGGCCTTTTTACGGTTCCTGGCCTTTTGCTGGCCTTTTGCTCACATGTTCTTTCCTGCGTTATCCCCTGATTCTGTGGATAACCGTATTACCGCCTTTGAGTGAGCTGATACCGCTCGCCGCAGCCGAACGACCGAGCGCAGCGAGTCAGTGAGCGAGGAAGCGGAAGAGCGCCCAATACGCAAACCGCCTCTCCCCGCGCGTTGGCCGATTCATTAATGCAGCTGGCACGACAGGTTTCCCGACTGGAAAGCGGGCAGTGAGCGCAACGCAATTAATGTGAGTTAGCTCACTCATTAGGCACCCCAGGCTTTACACTTTATGCTTCCGGCTCGTATGTTGTGTGGAATTGTGAGCGGATAACAATTTCACACAGGAAACAGCTATGACCATGATTACGCCAAGCGCGCAATTAACCCTCACTAAAGGGAACAAAAGCTGGAGCTCCACCGCGGTGGCGGCCGCCCCCTTCACCGAGGGCCTATTTCCCATGATTCCTTCATATTTGCATATACGATACAAGGCTGTTAGAGAGATAATTGGAATTAATTTGACTGTAAACACAAAGATATTAGTACAAAATACGTGACGTAGAAAGTAATAATTTCTTGGGTAGTTTGCAGTTTTAAAATTATGTTTTAAAATGGACTATCATATGCTTACCGTAACTTGAAAGTATTTCGATTTCTTGGCTTTATATATCTTGTGGAAAGGACGAAACACCGGTGTGCAGGTGAGTGATCCAAACGCCCGGCGGCAACCGAGCGTTCTGAACAAATCCAGATGGAGTTCTGAGGTCATTACTGGATCTATCAACAGGAGTCCAAGCGAGCTCTCGAACCCCAGAGTCCCGCTCAGAAGAACTCGTCAAGAAGGCGATAGAAGGCGATGCGCTGCGAATCGGGAGCGGCGATACCGTAAAGCACGAGGAAGCGGTCAGCCCATTCGCCGCCAAGCTCTTCAGCAATATCACGGGTAGCCAACGCTATGTCCTGATAGCGGTCCGCCACACCCAGCCGGCCACAGTCGATGAATCCAGAAAAGCGGCCATTTTCCACCATGATATTCGGCAAGCAGGCATCGCCATGGGTCACGACGAGATCCTCGCCGTCGGGCATGCGCGCCTTGAGCCTGGCGAACAGTTCGGCTGGCGCGAGCCCCTGATGCTCTTCGTCCAGATCATCCTGATCGACAAGACCGGCTTCCATCCGAGTACGTGCTCGCTCGATGCGATGTTTCGCTTGGTGGTCGAATGGGCAGGTAGCCGGATCAAGCGTATGCAGCCGCCGCATTGCATCAGCCATGATGGATACTTTCTCGGCAGGAGCAAGGTGAGATGACAGGAGATCCTGCCCCGGCACTTCGCCCAATAGCAGCCAGTCCCTTCCCGCTTCAGTGACAACGTCGAGCACAGCTGCGCAAGGAACGCCCGTCGTGGCCAGCCACGATAGCCGCGCTGCCTCGTCCTGCAGTTCATTCAGGGCACCGGACAGGTCGGTCTTGACAAAAAGAACCGGGCGCCCCTGCGCTGACAGCCGGAACACGGCGGCATCAGAGCAGCCGATTGTCTGTTGTGCCCAGTCATAGCCGAATAGCCTCTCCACCCAAGCGGCCGGAGAACCTGCGTGCAATCCATCTTGTTCAATCATGCGAAACGATCCTCATCCTGTCTCTTGATCAGATCTTGATCCCCTGCGCCATCAGATCCTTGGCGGCAAGAAAGCCATCCAGTTTACTTTGCAGGGCTTCCCAACCTTACCAGAGGGCGCCCCAGCTGGCAATTCCGACGGATCAAGAACCTGCTGACGTTTTAGAGCTAGAAATAGCAAGTTAAAATAAGGCTAGTCCGTTATCAACTTGAAAAAGTGGGACCGAGTCGGTCCTTTTTTTGAATTCGATATCAAGCTTATCGATACCGTCGACCTCGAGGGGGGGCCCGGTACCCAATTCGCCCTATAGTGAGTCGTATTACGCGCGCTCACTGGCCGTCGTTTTACAACGTCGTGACTGGGAAAACCCTGGCGTTACCCAACTTAATCGCCTTGCAGCACATCCCCCTTTCGCCAGCTGGCGTAATAGCGAAGAGGCCCGCACCGATCGCCCTTCCCAACAGTTGCGCAGCCTGAATGGCGAATGGGACGCGCCCTGTAGCGGCGCATTAAGCGCGGCGGGTGTGGTGGTTACGCGCAGCGTGACCGCTACACTTGCCAGCGCCCTAGCGCCCGCTCCTTTCGCTTTCTTCCCTTCCTTTCTCGCCACGTTCGCCGGCTTTCCCCGTCAAGCTCTAAATCGGGGGCTCCCTTTAGGGTTCCGATTTAGTGCTTTACGGCACCTCGACCCCAAAAAACTTGATTAGGGTGATGGTTCACGTAGTGGGCCATCGCCCTGATAGACGGTTTTTCGCCCTTTGACGTTGGAGTCCACGTTCTTTAATAGTGGACTCTTGTTCCAAACTGGAACAACACTCAACCCTATCTCGGTCTATTCTTTTGATTTATAAGGGATTTTGCCGATTTCGGCCTATTGGTTAAAAAATGAGCTGATTTAACAAAAATTTAACGCGAATTTTAACAAAATATTAACGCTTACAATTTAG
